# A Genetically Encoded Reporter for Diffusion Weighted Magnetic Resonance Imaging

**DOI:** 10.1101/037515

**Authors:** Arnab Mukherjee, Di Wu, Hunter C. Davis, Mikhail G. Shapiro

## Abstract

The ability to monitor gene expression in intact, optically opaque animals is important for a multitude of applications including longitudinal imaging of transgene expression and long term tracking of cell based therapeutics. Magnetic resonance imaging (MRI) could enable such monitoring with high spatial and temporal resolution. However, existing MRI reporter genes, based primarily on metal-binding proteins or chemical exchange saturation transfer probes, are limited by their reliance on metal ions or relatively low sensitivity. In this work, we introduce a new class of genetically encoded reporters for MRI that work by altering water diffusivity. We show that overexpression of the human water channel aquaporin 1 (AQP1) produces robust contrast in diffusion weighted MRI by increasing effective water diffusivity in tissues by over 100% without affecting cell viability or morphology. Low levels of AQP1 expression (˜1 μM), or mixed populations comprising as few as 10% AQP1-expressing cells, produce sufficient contrast to be observed by MRI. We demonstrate the utility of AQP1 *in vivo* by imaging gene expression in intracranial tumor xenografts. Overall, our results establish AQP1 as a new, metal-free, nontoxic and sensitive genetically encoded reporter for diffusion weighted MRI.

## INTRODUCTION

The ability to image gene expression within the context of living mammalian organisms is critical for basic biological studies and the development of cellular and genetic therapeutics. However, most genetically encoded reporters, based on fluorescent and luminescent proteins^1-3^ have limited utility in this context due to the poor penetration of light into deep tissues^1,2,4,5^. In contrast to optical techniques, magnetic resonance imaging (MRI) enables the acquisition of *in vivo* images with excellent depth penetration and high spatial and temporal resolution. Consequently, there is intense interest in the development of genetically encoded reporters for MRI^6-28^. Previous efforts to develop such reporters have focused primarily on two classes of proteins. In one class, metalloproteins and metal ion transporters are overexpressed to enrich the paramagnetic content of cells, thereby enhancing nuclear relaxation rates and producing contrast in T_1_ or T_2_-weighted MRI^9,12-19,27-29^. In the second strategy, proteins with large numbers of basic amino acids are used to generate contrast through chemical exchange saturation transfer (CEST) between protein-bound and aqueous protons^6,8,21,22,25,30^. Each of these pioneering approaches has significant limitations. Metal-based reporters can be hindered by metal ion bioavailability and toxicity^31-35^, while CEST reporters tend to require high expression levels to achieve observable contrast^6,21,22,30^. Hence, a major need exists for new MRI reporter genes that do not require metals and can be detected at low levels of expression.

Here, we introduce an entirely new class of non-metallic MRI reporter genes that work by modulating water diffusivity across cell membranes. Diffusion weighted imaging (DWI) is a well-established MRI modality used in a wide range of applications from basic biophysical studies to the diagnosis of diseases such as stroke^36-40^. Diffusion weighting is commonly achieved by applying a pair of pulsed magnetic field gradients, which dephase proton spins proportionally to their diffusion distance in the time interval between gradient applications^41-43^. Accordingly, tissue regions characterized by rapid water diffusion have reduced signal intensity compared to regions with restricted water mobility. In biological tissues, the effective diffusion coefficient of water depends on several parameters including the local diffusivity in intracellular and extracellular compartments, the relative volume fraction occupied by cells, and the diffusion of water across the plasma membrane^44-49^. Noting the strong influence of the last factor^44,50^, we hypothesized that facilitating the transmembrane diffusion of water by overexpressing water-permeable channels would result in enhanced contrast in DWI. Towards this end, aquaporins are a highly conserved family of tetrameric integral membrane proteins that mediate the selective exchange of water molecules across the plasma membrane in a wide range of cell types including erythrocytes, astrocytes, and kidney cells^51-53^. Previously, endogenous aquaporin expression has been correlated with water diffusivity and DWI signals in several disease states^52,54,55^. However, to the best of our knowledge, aquaporins have not hitherto been described as MRI reporter genes. In this work, we introduce human aquaporin 1 (AQP1) as a new genetically encoded reporter for diffusion weighted MRI. This reporter gene requires no metals, is nontoxic in a wide range of cells, produces contrast orthogonal to paramagnetic and CEST reporters and is detectable when expressed at low levels and in small subsets of cells. We characterize the imaging performance and mechanisms of AQP1 through live cell experiments and Monte Carlo models, and demonstrate its utility by imaging tumor gene expression *in vivo.*

## RESULTS

### Aquaporins serve as reporter genes for diffusion weighted MRI

To evaluate AQP1 as a genetically encoded reporter for diffusion weighted MRI (**Figure 1a**),we used lentiviral transfection to generate several mammalian cell lines stably overexpressing this channel or green fluorescent protein (GFP). Pellets of AQP1-expressing and GFP-expressing CHO, U87 glioblastoma, and Neuro2A neuroblastoma cells were then imaged using DWI. A key parameter in diffusion weighted pulse sequences is the effective diffusion time, A eff, corresponding to the time interval between dephasing and rephasing gradient pulses^36,37,43,44,46,56^. Long Δ_eff_ times are important for probing the effects of water exchange between intracellular and extracellular pools because longer times allow a larger proportion of cytoplasmic molecules to interact with the cell membrane and experience the effects of restriction and exchange^36,37,46,49^. Correspondingly, Monte Carlo simulations of a packed cellular lattice suggested that the effects of an aquaporin-mediated increase in water diffusion would be most pronounced at Δ_eff_ > 100 ms **(Figure S1)**. To access these longer diffusion times, we used a stimulated echo DWI sequence, in which net magnetization is stored along the longitudinal axis in the interval between the diffusion gradients, and is thereby limited by T_1_ relaxation, rather than the typically shorter T_2_ relaxation limit of the more widely used spin echo DWI^44,56,57^.

**Figure 1:**
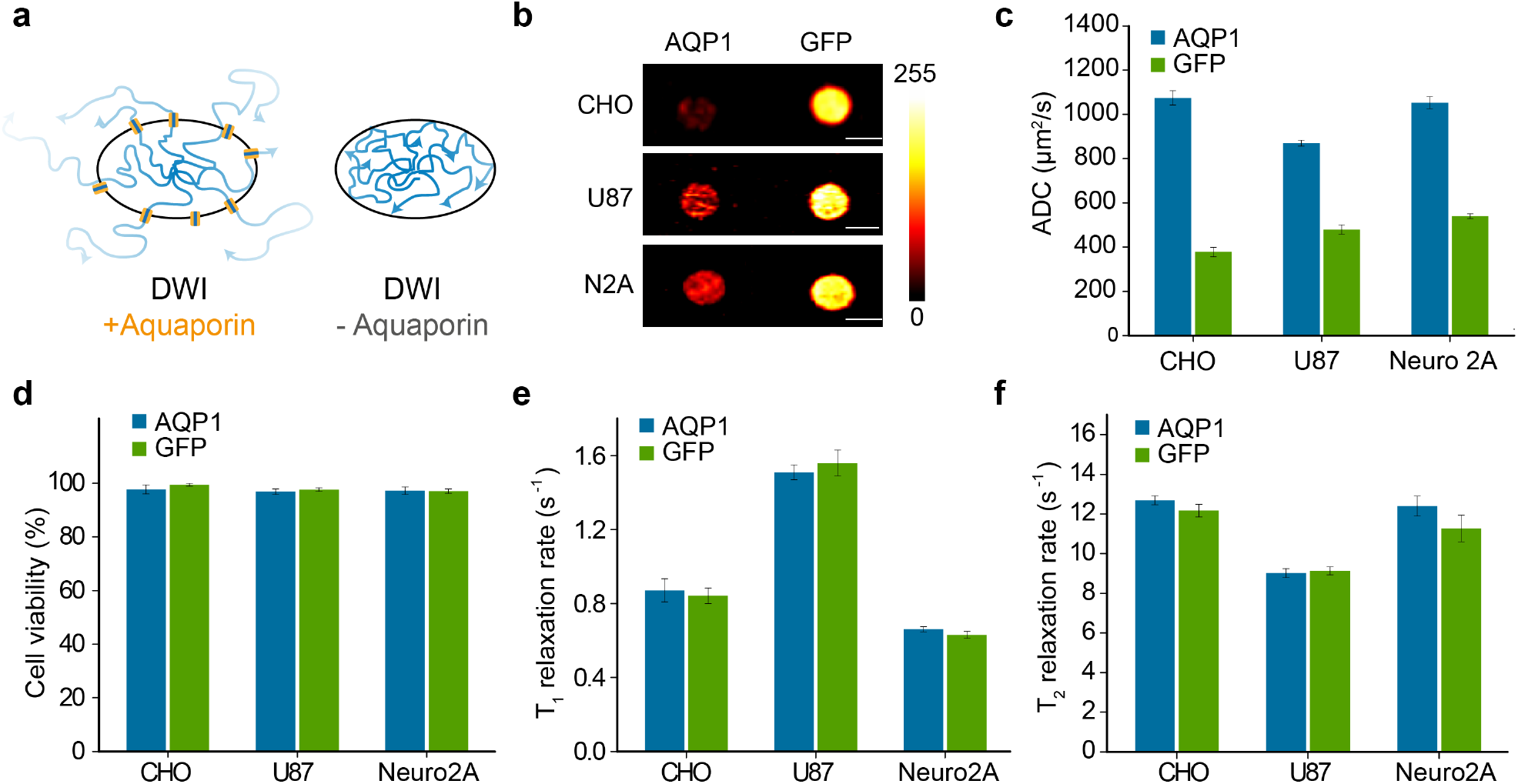
AQP1 functions as a genetically encoded reporter for diffusion weighted MRI. (a) Illustration of the impact of aquaporin expression on water diffusion across the cell membrane and the resulting decrease in diffusion-weighted signal intensity. (b) Diffusion weighted images of CHO, U87, and Neuro2A cell pellets expressing AQP1 or GFP. The images were acquired at Δ_eff_ = 298 ms with a b-value of 2089 s/mm^2^ (CHO), 1000 s/mm^2^ (U87), and 800 s/mm^2^ (Neuro2A). Scale bar, 3 mm. (c) ADC of water in CHO, U87, and Neuro2A cells expressing AQP1 relative to GFP controls, measured at Δ_eff_ = 398 ms. Transgene expression in CHO cells was induced with 1 pg/μL doxycycline, while U87 and Neuro2A cells express AQP1 from a constitutive promoter. (d) Cell viability upon AQP1 or GFP expression determined with ethidium homodimer-1 assay. (e) Longitudinal (T_1_) and (f) transverse (T_2_) relaxation rates in cells expressing AQP1 or GFP. Error bars represent standard error of mean (SEM) for 4 biological replicates.

Pellets of AQP1-expressing cells appeared much darker in diffusion-weighted images than GFP controls for all cell types (**Figure 1b**), corresponding to dramatic increases in their apparent diffusion coefficients (ADC, **Figure 1c**). Measured with Δ_eff_ = 398 ms, AQPl-expressing CHO, U87 and Neuro2A cells showed 187 ± 4%, 82 ± 5% and 95 ± 3% increases in ADC, respectively, compared to GFP controls. The relative increase in ADC is more pronounced at Δ_eff_ = 398 ms compared to Δ_eff_ = 18 ms **(Figure S2)**, consistent with AQP1 expression facilitating the exchange of water across the cell membrane. The larger change in ADC in CHO cells compared to Neuro2A and U87 is a likely consequence of the stronger, inducible promoter used to drive expression in CHO cells compared to the relatively weaker constitutively active promoter used in the two other cell types. Importantly, AQP1 overexpression was nontoxic in all cell lines, as determined by staining with ethidium homodimer 1, which identifies cells with compromised membrane integrity (**Figure 1d**). In addition, no changes in cell size or morphology were observed under phase contrast microscopy as a result of AQP1 expression.

To establish orthogonality to paramagnetic reporters, we measured the T_1_ and T_2_ relaxation rates of cells expressing AQP1. Overexpression of this protein was not seen to affect T_1_ or T_2_ relaxation (**Figure 1, e-f**), suggesting that AQP1 could be used in combination with genetically encoded T_1_ or T_2_ contrast agents for multiplexed imaging. We note that we were also able to obtain a significant increase in ADC by transfecting cells with another human aquaporin, AQP4 **(Figure S3)**. However, the percentage increase in ADC for the AQP4 expressing cells (44 ± 6% in CHO cells) was smaller compared to the AQP1 expressing cells. Therefore, we chose to focus on AQP1 for the remainder of the work.

### AQP1 is a sensitive reporter of gene expression across a large dynamic range

Next, we sought to establish whether AQP1 can be used to report on varying degrees of gene expression, particularly at low levels of expression. Our Monte Carlo simulations suggested that ADC values are sensitive to a broad range of cell membrane permeabilities **(Figure S1b)**, providing AQP1 with significant dynamic range. To realize this experimentally, we expressed AQP1 in a dose-dependent fashion by supplementing CHO cells with varying concentrations of doxycycline and measured corresponding values of ADC (**Figure 2, a-b**). AQP1 expression was also quantified via western blotting (**Figure 2c**). Significant changes in ADC and DWI contrast (53% and 29% respectively) were observed with very low levels of doxycycline induction (0.01 μg/mL), which corresponded to membrane AQP1 expression below the chemiluminescence detection limit of our western blot. At the lowest blotting-detectable level of AQP1 expression, corresponding to an AQP1 concentration of 1.06 ± 0.19 μM (induced with 0.1 μg/mL doxycycline, N = 2), cells showed a 164 ± 5% increase in ADC relative to controls. Since substantial DWI contrast is also observed at 10-fold lower induction levels, we expect that the actual detection limit for AQP1 expression is significantly below 1 μM. This large dynamic range will facilitate the use of AQP1 as a reporter gene in a variety of biomedical applications.

**Figure 2:**
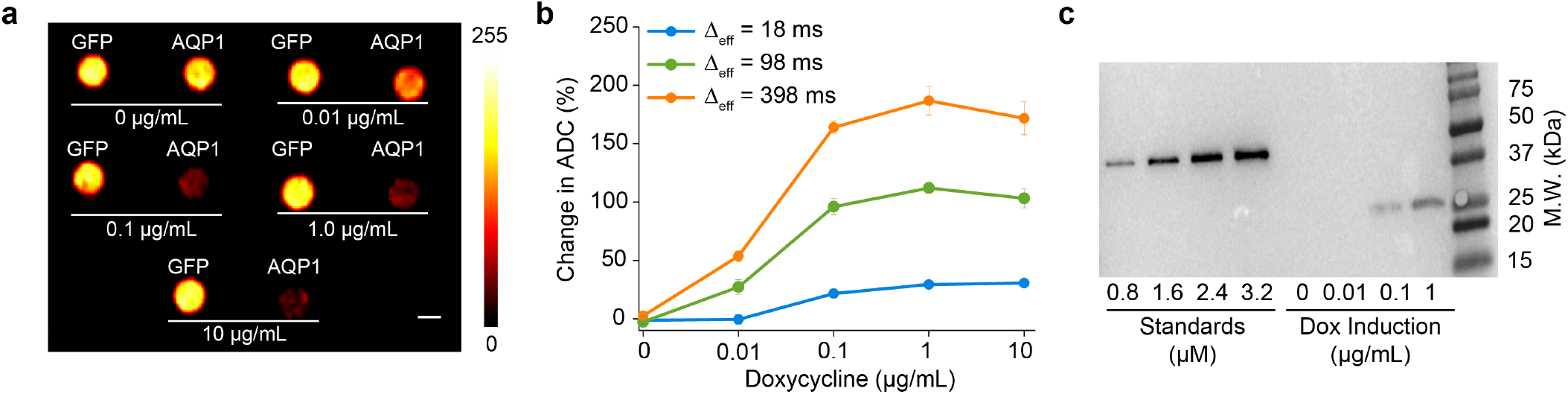
AQP1 reports gene expression over a large dynamic range. (a) Diffusion weighted images (acquired at Δ_eff_ = 398 ms, b = 2089 s/mm^2^) of CHO cells expressing AQP1 or GFP (control) and treated with varying doses of doxycycline to induce transgene expression. Scale bar indicates 3 mm. (b) Percent change in ADC of water in AQP1-expressing CHO cells (relative to control cells expressing GFP) as a function of doxycycline concentration, measured at different diffusion times. Error bars represent SEM for 4 biological replicates. (c) Representative western blot used to quantify AQP1 expression in CHO cells using FLAG-tagged bacterial alkaline phosphatase at the indicated concentrations as a calibration standard. AQP1 levels were estimated using membrane fractions isolated from AQP1 expressing CHO cells respectively induced using 0, 0.01, 0.1, and 1 μg/mL doxycycline for 48 hours. AQP1 levels corresponding to 0.01 μg/mL doxycycline are below the blottingdetectable limit.

### AQP1 expression is observable in small subsets of cells within a mixed population

The ability to specifically detect small numbers of genetically labeled cells in a population of unlabeled cells would enable the use of genetically encoded reporters in applications such as *in vivo* tracking of cell based therapeutics^16,58,59^. Having shown that AQP1 can appreciably increase water diffusion even at low levels of expression (**Figure 2**), we tested whether apparent water diffusion could be significantly increased if AQP1 expression was restricted to a small subset of cells in a mixed population. In general, the relationship between expressing fraction and ADC is expected to be nonlinear, since in small-fraction scenarios, cells expressing AQP1 would be surrounded mostly by cells without enhanced water permeability, and the impact of AQP1 expression would therefore be diminished (**Figure 3a**). However, our Monte Carlo simulations predicted that even in this scenario, expressing fractions as small as 10% could be sufficient to increase the overall ADC in heterogeneous cell populations, particularly at long Δ_eff_ times (**Figures 3b, S1c)**. To verify this experimentally, we measured ADC in mixed populations comprising AQP1 expressing CHO cells and GFP expressing control cells in varying proportions. Strikingly, diffusion measurements revealed a significant increase in ADC in cell populations comprising 10% AQP1 expressing cells (21.44 ± 5.21% relative to GFP expressing cells, measured at Δ_eff_ = 398 ms, *P = 0.04, n = 4*, **Figure 3c-d**). Furthermore, under optimal imaging conditions we were able to observe 12.87% and 19.58% decreases in diffusion weighted image intensity for 5% and 10% AQP1 cell populations, respectively, relative to homogeneous GFP controls (**Figure 3c, inset)**. This data suggests that, contrary to initial intuition, diffusional reporter genes such as AQP1 are suitable for imaging gene expression in heterogeneous or infiltrating cell populations.

**Figure 3:**
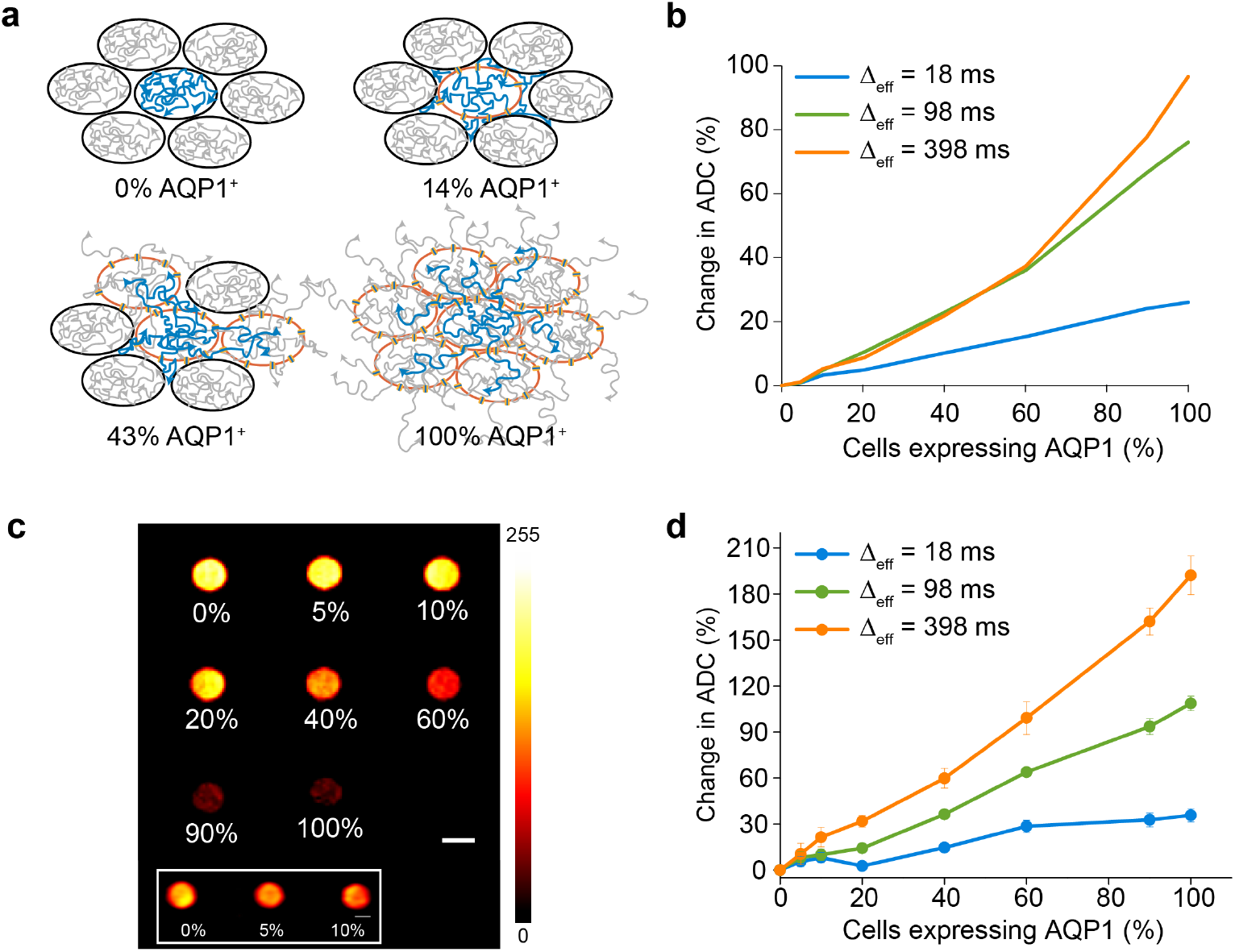
AQP1 expression is observable in mixed cell populations. (a) Illustration of the effect of an increasing fraction of AQP1-labeled cells in a tissue on the overall diffusivity of water. (b) Monte Carlo simulations of percent increase in ADC as a function of the fraction of cells expressing AQP1 in a mixed population. (c) Diffusion weighted MRI (acquired at Δ_eff_ = 198 ms, b = 2334 s/mm^2^) of cells comprising AQP1-labeled cells mixed with GFP-labeled control cells in varying proportions. (Inset) Mixed populations consisting of 0, 5, and 10% AQP1 expressing cells independently imaged using optimal parameters (Δ_eff_ = 398 ms, b = 8000 s/mm^2^) to maximize contrast for the low AQP1 fraction scenario. In order to reduce image noise and improve visual clarity, the image was smoothed using a low pass Gaussian filter, implemented in ImageJ. Scale bar represents 3 mm. (d) Experimental percent change in ADC in mixed AQP1/GFP cell pellets as a function of the fraction of AQP1 expressing cells. Error bars represent SEM for 4 biological replicates.

### AQP1 enables the imaging of gene expression in intracranial tumor xenografts

To demonstrate the ability of AQP1 to report gene expression *in vivo*, we stereotaxically implanted AQP1 and GFP-transfected CHO cells in the right and left striatum of 5-7 week old NOD/SCID/gamma nude mice. Tumors were allowed to develop for a period of 5 days, following which AQP1 and GFP expression was induced via intraperitoneal injection of 75 μg doxycycline. Mice were imaged using diffusion weighted MRI before and 48 hours after doxycycline injection. Our experimental protocol is outlined in **Figure 4a.** As expected, AQP1-expressing tumors are readily distinguishable from contralateral GFP-expressing tumors in diffusion weighted images acquired after induction (**Figure 4b**), with the average signal intensity in AQP1 tumors decreasing by 40.72% after doxycycline injection compared to GFP controls (pairwise t-test, *P = 0.01, n = 5)* (**Figure 4c, S5)** The use of an intermediate-length Δ_eff_ (98 ms), for these *in vivo* experiments provided the optimal balance of AQP1-dependent contrast and acquisition times. AQP1 and GFP expression in the bilateral tumors was confirmed by fluorescence imaging of fixed brain tissue slices (**Figure 4d**). The AQP1 tumor shows weak green fluorescence based on the presence of an IRES-driven GFP gene in the construct. The ability of AQP1 to produce robust induction-dependent MRI contrast in tumor xenograts suggests that this reporter gene could be useful for longitudinal imaging of gene expression in intact animals.

**Figure 4:**
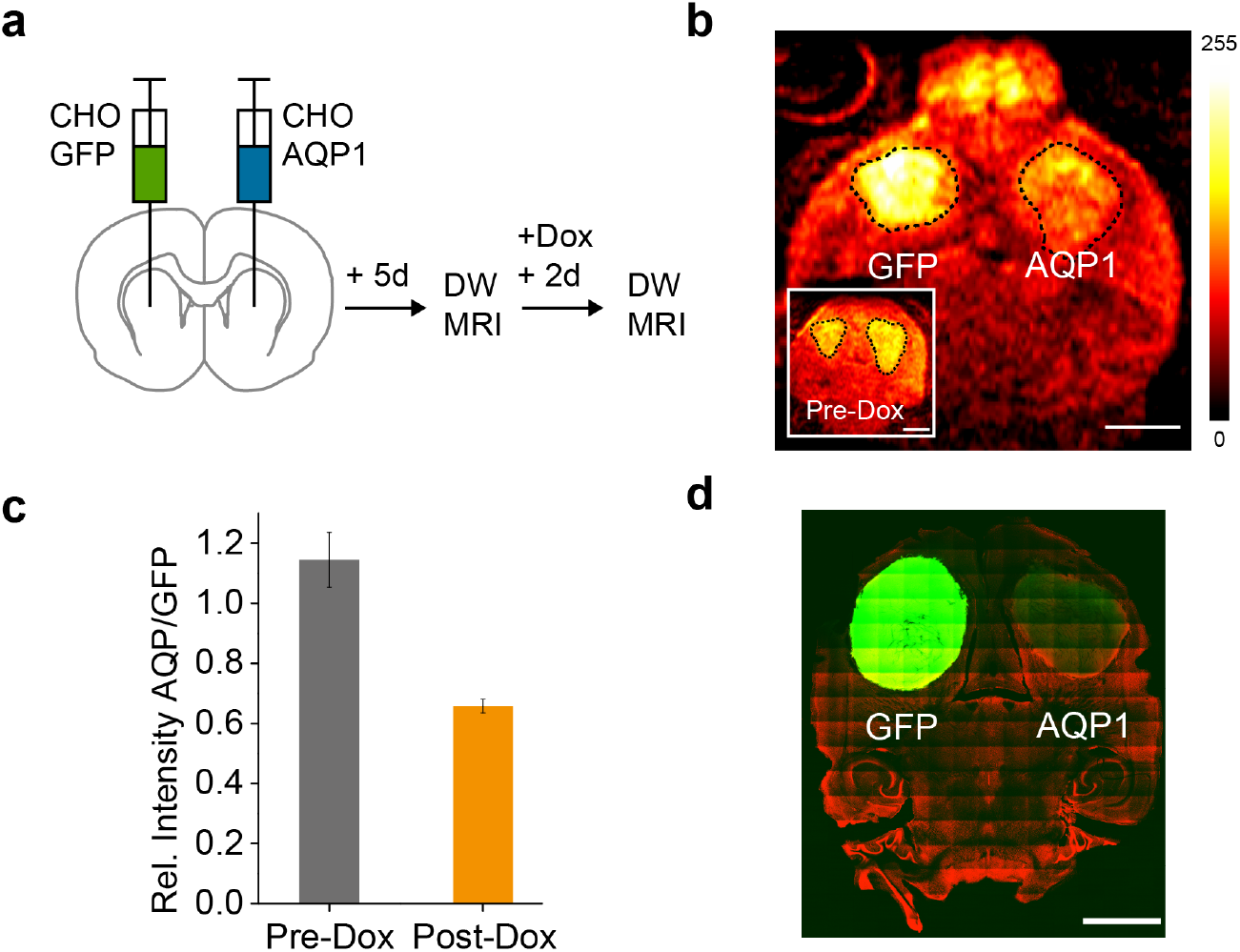
AQP1 as a reporter for imaging intracranial tumor xenografts. (a) Schematic outline of experimental approach for establishing bilateral tumors in the striatum, inducing transgene expression, and diffusion weighted MRI. (b) Representative diffusion weighted image of a horizontal section of a mouse brain with a bilateral tumor xenograft, 48 hours following doxycycline injection. (Inset) Diffusion weighted image of the same mouse acquired 48 hours prior to doxycycline injection, depicting similar intensities in the AQP1 and GFP-labeled tumors. Diffusion weighted images were acquired at Δ_eff_ = 98 ms and b-value = 1000 s/mm^2^. Dashed lines indicate the tumor ROI(s). Scale bar represents 3 mm. (c) Normalized DWI intensity of AQP1-expressing tumors before and after AQP1 induction via intraperitoneal injection of doxycycline. Error bars represent SEM for 5 biological replicates. (d) Confocal image of a 100 μm histological section of a mouse brain. Transgene expression in the tumors is indicated by bright GFP fluorescence in the left tumor and weak fluorescence in the right tumor owing to diminished translation from the IRES sequence. Nuclei (shown in red) are counterstained using TO-PRO iodide. Scale bar is 3 mm.

## DISCUSSION

Our results establish aquaporins, and specifically AQP1, as the first genetically encoded reporter for diffusion weighted MRI. AQP1-dependent contrast is readily observed in cell cultures, including cells known to have higher levels of endogenous aquaporins *(e.g.*, U87 glioblastoma cells^60^) as well as *in vivo* tumor xenografts. In contrast to metal-based agents, AQP1 does not require metal ions or other exogenous factors. Importantly, low levels of AQP1 expression (<1 μM) were found to be sufficient to enhance ADC and produce contrast in diffusion weighted MRI, placing it among the most sensitive MRI reporters. Moreover, AQP1-mediated increase in ADC was reliably detected even when the fraction of AQP1 expressing cells comprised only 10% of the overall population. In this regard, AQP1 can potentially fulfil the need for a sensitive and nontoxic reporter for specifically labeling and tracking immune and stem cell-based therapeutics for preclinical and clinical applications. Furthermore, AQP1 expression does not affect transverse and longitudinal relaxation rates in cells, which opens the door to multiplexed imaging of gene expression by combining AQP1 with T_1_, T_2_, or CEST reporters. In addition, we anticipate that aquaporins as a class of proteins will readily lend themselves to molecular engineering of variants with improved or stimulus-gated permeability to enable functional imaging of biologically relevant markers.

Given the ubiquity of DWI and stimulated echo pulse sequences, the imaging of aquaporin-based reporters can be implemented immediately by laboratories with standard MRI equipment. A potential limitation of these simple pulse sequences is the requirement for long diffusion times to develop membrane permeability-dependent DWI contrast and the accompanying loss of signal due to T1 relaxation. Consequently, imaging under these conditions typically requires multiple excitations for signal averaging, which impacts the temporal resolution achievable using aquaporins. This limitation could potentially be overcome with alternative pulse sequences specifically designed to produce contrast based on transmembrane water exchange^57^, the development of which will be further stimulated by the present work. Overall, the high performance, biocompatibility and engineering capacity of aquaporin-based reporter genes will enable a broad array of applications ranging from basic biological studies to long-term tracking of cell therapies and imaging the expression of endogenous and therapeutic genes.

## MATERIALS AND METHODS

Detailed methods are available as supporting information.

### Construction of aquaporin and GFP expressing stable cell lines

Human aquaporins 1 and 4 were ordered from OriGene (Rockville, MD) and subcloned in a lentiviral vector downstream of a constitutive CMV or doxycycline regulated CMVTight promoter (Clontech) and an N-terminal FLAG tag. EGFP was fused downstream of AQP via an IRES sequence. Cloning was accomplished using Gibson assembly. Stable, polyclonal CHO, CHO rtTA, Neuro 2A, and U87 cell lines were derived by lentiviral transfection at high MOI(s) to achieve near complete transfection efficiency. Control cell lines were generated in the same way, and they express EGFP from a constitutive full length CMV or a doxycycline regulated CMVTight promoter. AQP1 expression was detected and quantified using western blotting.

### Diffusion weighted MRI of aquaporin expression in cell pellets

For diffusion weighted MRI of cells, CHO, U87, or Neuro 2A cells were grown for 48 hours, trypsinized, resuspended in 100 μL PBS, and centrifuged at 500 x g for 5 minutes in 0.2 mL PCR tubes to produce a compact pellet. Subsequently, the tubes were loaded in wells molded in a 1% agarose phantom and imaged using a Bruker 7T horizontal bore MRI scanner equipped with a 7.2 cm diameter bore transceiver coil for RF excitation and detection. Diffusion weighted images were acquired on a 1.5 or 2 mm thick horizontal slice through the cell pellets using a stimulated echo DWI sequence with the following parameters: echo time, T_E_ = 24.5 ms, repetition time, T_R_ = 2 s, number of excitations = 1 — 3, gradient duration, δ = 7 ms, matrix size = 256 x 256, FOV = 3.5 x 6.5 cm^2^. The gradient interval (Δ) was varied from 20 to 500 ms to generate effective diffusion times (Δ_eff_ = Δ- δ/3) of 18—498 ms in each experiment. Single axis diffusion gradients were applied and gradient strength was varied to generate b-values in the range 0—800 s/mm^2^ For each value of Δ_eff_, ADC was calculated from the slope the logarithmic decay in MRI signal intensity versus b-value — that is, ***ln(S/So) = −b •*** *ADC*, where *S* and *S_0_* denote MR signal intensities in the presence and absence of diffusion gradients. All images were analyzed using custom macros in ImageJ (NIH). The maximum and minimum values of a linear 8-bit color scale were chosen to facilitate the visualization of the relevant contrast in each figure. Least squares regression fitting was performed using OriginLab.

### Mouse xenograft model

To prepare cells for intracranial tumor implantation, AQP1 and GFP expressing CHO-rtTA cells were grown for 48 hours, trypsinized, centrifuged at 500 x g for 10 minutes, and resuspended in 100 μL serum-free Dulbecco’s Modified Eagle Medium. Female NOD/SCID/gamma mice between 5 and 7 weeks of age (Jackson Laboratory, Bar Harbor, ME) were anaesthetized with 2.5% isoflurane-air mixture and ˜10^5^ AQP1 expressing CHO cells were injected into the right striatum guided by a small-animal stereotaxic frame (Kopf instruments, Tujunga, CA). Coordinates of the injection sites with respect to bregma were: 1 mm anterior, 2 mm lateral, and 1 —3 mm beneath the surface of the skull. Control GFP expressing CHO cells were implanted in the left striatum of the same animal. Tumor growth and gene expression were confirmed by histology. All animal surgical protocols were approved by the Institutional Animal Care and Use Committee of the California Institute of Technology.

### Diffusion weighted MRI of mouse xenografts

Diffusion weighted imaging of mouse xenografts was performed using a Bruker 7T horizontal bore MRI scanner. RF excitation was delivered by a 7.2 cm diameter bore volume coil and detection was achieved using a 3 cm diameter surface coil. Mice were lightly anaesthetized using 2% isoflurane-air mixture and placed on a dedicated animal bed with the surface coil positioned proximal to the head of the mouse. Anaesthesia was maintained over the course of imaging using 1-1.5% isoflurane. Warm air was circulated in the bore of the MRI scanner to maintain body temperature at 30°C. Respiration rate was maintained at 20-30 breaths per minute and temperature and respiration rate were continuously monitored during the imaging session using a pressure pad/respiration transducer (Biopac Systems) and a fiber optic rectal temperature sensor (Neoptix). Tumor formation was confirmed by acquiring diffusion weighted images 5 days following xenograft implantation after which mice were intraperitoneally injected with 75 μg doxycycline to induce expression of AQP1 and GFP in the tumors. A second set of diffusion weighted images was acquired 48 hours following doxycycline injection. Preliminary diffusion weighted images to locate the tumors were first acquired on horizontal slices using a 3D echo planar imaging (EPI) stimulated echo DWI sequence with the following parameters: T_R_ = 2.5 or 3 s, T_E_ = 25.7 ms, δ= 7 ms, Δ = 100 ms, b = 1000 s/mm^2^, number of excitations = 9, matrix size = 16 x 128 x 128, FOV = 1.59 x 1.29 x 0.74 cm^3^. After identifying the tumor bearing slice, 2D EPI diffusion weighted images were acquired at the slice using similar parameters but with a slice thickness of 2 mm, T_R_ = 5 s, number of excitations = 144—256. All animal imaging protocols were approved by the Institutional Animal Care and Use Committee of the California Institute of Technology.

## ACKNOWLEDGEMENTS

We acknowledge George Lu, Pradeep Ramesh, Russell Jacobs, Xiaowei Zhang and Michael Tyszka for useful discussions, and Philip Petersen for experimental contributions. AM acknowledges financial support from the James G. Boswell foundation. DW was supported by a Medical Engineering Amgen Fellowship. The work was funded by a grant from the Dana Foundation and the Burroughs Wellcome Career Award at the Scientific Interface

